# High-resolution fleezers reveal duplex opening and stepwise assembly by an oligomer of the DEAD-box helicase Ded1p

**DOI:** 10.1101/2024.02.29.582829

**Authors:** Eric M. Patrick, Rajeev Yadav, Kasun Senanayake, Kyle Cotter, Andrea A. Putnam, Eckhard Jankowsky, Matthew J. Comstock

## Abstract

DEAD-box RNA helicases are ubiquitous in all domains of life where they bind and remodel RNA and RNA-protein complexes. DEAD-box helicases unwind RNA duplexes by local opening of helical regions without directional movement through the duplexes and some of these enzymes, including Ded1p from Saccharomyces cerevisiae, oligomerize to effectively unwind RNA duplexes. Whether and how DEAD-box helicases coordinate oligomerization and unwinding is not known and it is unclear how many base pairs are actively opened. Using high-resolution optical tweezers and fluorescence, we reveal a highly dynamic and stochastic process of multiple Ded1p protomers assembling on and unwinding an RNA duplex. One Ded1p protomer binds to a duplex-adjacent ssRNA tail and promotes binding and subsequent unwinding of the duplex by additional Ded1p protomers in 4-6 bp steps. The data also reveal rapid duplex unwinding and rezipping linked with binding and dissociation of individual protomers and coordinated with the ATP hydrolysis cycle.

## INTRODUCTION

RNA helicases are critical in all steps of RNA metabolism^1,2^. Mutations and dysregulation of RNA helicases have been linked to numerous diseases and there is considerable interest in understanding molecular functions, roles in diseases and means to therapeutically target these enzymes^3^. All RNA helicases utilize nucleoside triphosphates to bind and remodel RNA, RNA-protein complexes, or both^3^. RNA helicases are classified into the DEAD-box, DEAH/RHA, Ski2-like, Upf1-like and RIG-I families, based on sequence and structural features^3^. Except for the RIG-I-like family, most, if not all RNA helicases unwind RNA duplexes *in vitro*, provided suitable substrates are used^4^. RNA unwinding by DEAD-box helicases is not based on translocation along the RNA, in contrast to canonical DNA helicases and other RNA helicases^4^. Instead of translocating, DEAD-box helicases load directly to the duplex and pry apart a limited number of base pairs in an ATP-dependent fashion, while the remaining base pairs dissociate non-enzymatically^5–13^. Loading onto the duplex is facilitated by unpaired RNA proximal to the duplex^8,12^. Loading can be accomplished by unpaired regions of either polarity, in contrast to canonical translocating helicases, for which unpaired regions must match translocation directionality^2,14–18^.

Some DEAD-box helicases form oligomers to efficiently unwind RNA duplexes^19–22^, including Ded1p from *Saccharomyces cerevisiae*, and its human ortholog DDX3X^19^. Both Ded1p and DDX3X are involved in translation initiation^23–29^ and have also been implicated in other aspects of RNA metabolism^22,30–39^. Mutations and dysregulation of DDX3X has been linked to various diseases, including medulloblastoma, several other cancers and the eponymous DDX3 syndrome, a neurodevelopmental disorder^3,40–44^.

Despite the considerable interest in Ded1p, DDX3X and DEAD-box helicase mechanisms in general, and despite the considerable body of work on these enzymes that has accumulated over more than three decades, including recent single molecule work,^45^ several fundamental mechanistic aspects of DEAD-box helicase function are not known. Firstly, how many base pairs DEAD-box helicases actively open during duplex unwinding has not been measured. This number is a key parameter for helicase activity, as it dictates unwinding efficiency. Secondly, it is not clear how oligomeric DEAD-box helicases coordinate oligomerization, substrate binding, unwinding, and the ATP hydrolysis cycle. While ensemble investigations have shown that a triplet of Ded1p protomers are involved in RNA duplex unwinding at high Ded1p concentration, it is unclear to what extent proteins assemble in solution vs on the RNA substrate or to what extent individual proteins contribute to RNA unwinding vs reaction scaffolding^19^. Insight into these unsolved mechanistic questions for Ded1p and DDX3X is of high value for the development of therapeutics targeting DDX3X.

To address these questions, we used high-resolution optical tweezers combined with single molecule fluorescence (fleezers) to directly measure duplex opening for Ded1p, detailed assembly of the Ded1p oligomer and coupling of unwinding activity and oligomerization, all at the single molecule level. Our data show that a single Ded1p protomer binds to the duplex-adjacent unpaired RNA, which then promotes binding of additional Ded1p protomers to the duplex. We observe active opening of the duplex by Ded1p in 4-6 bp increments. In addition, our data reveal rapid duplex unwinding and rezipping linked with binding and dissociation of individual protomers and coordination of strand opening and Ded1p protomer binding and dissociation with the ATP hydrolysis cycle. Unwinding is highly dynamic with individual protomers binding and dissociating from the RNA substrate. Stability is enhanced by the binding of neighboring promoters, leading to the strong enhancement in unwinding activity with increasing protein concentration and oligomerization on the RNA substrate.

## RESULTS

### Ded1p unwinds duplex RNA in multiple reversible steps

We investigated Ded1p unwinding a typical RNA substrate at the single molecule level using high-resolution optical tweezers. **Figure 1a** shows a schematic of the tweezers assay where a 16 bp RNA hairpin with adjacent 25 nt 3’ overhang ligated between a pair of ∼1.5 kb dsDNA linkers (**Supplementary Fig. 1)** is held in between and monitored by a pair of trapped microspheres. The substrate was as previous model substrates where the overhang length was the minimum for promoting enhanced unwinding at increased Ded1p concentrations^9,12,19,46^. Single tether molecules were formed and tested in a ‘blank’ solution (contained ATP but no Ded1p) then transferred to an adjacent solution containing Ded1p. We performed a series of experiments increasing the Ded1p concentration from below to above the expected trimerization concentration (as suggested by previous ensemble experiments^9,19^). **Figure 1b** and **c** show example unwinding measurements at the extremes of low (100 nM, **Fig. 1b**) vs high (500 nM, **Fig. 1c**) Ded1p concentration. For hairpin unwinding tweezers assays such as these, unwinding and rezipping of the RNA hairpin is observed as increases and decreases respectively in signal (resulting from increases and decreases in the tether extension and calibrated to the corresponding number of RNA hairpin bp unwound; see Methods for details). Unwinding required ATP and the rate constants for unwinding and rezipping were ATP-concentration dependent (full discussion below). At low Ded1p concentration, we observed repeated partial unwinding of the RNA hairpin (complete unwinding corresponds to a 16 to 18 bp signal) followed by rapid rezipping. Typically, only a single approximately 5 to 6 bp unwinding step was observed followed quickly (∼0.25 sec) by reversal to fully zipped. Less often a second 5 to 6 bp unwinding step followed the first unwinding step. In contrast at high Ded1p concentration, the RNA hairpin was completely unwound rapidly, reversibly, and often (**Fig. 1c**). Two similar successive 5 to 6 bp unwinding steps were observed followed by smaller approximately 2 to 3 bp steps. Partial and complete rezipping steps were observed often. The complete unwinding reaction was so robust that repeated cycles of unwinding and rezipping continued for the entire tether lifetime often lasting tens of minutes and producing thousands of successive unwinding reactions. This robustness prompted the foundational question of whether this repeated activity was carried out by a single Ded1p complex anchored to the RNA hairpin or if fresh Ded1p protein bind and dissociate from solution. Therefore, we performed experiments where individual tethers were transferred between adjacent solutions with or without Ded1p (**Fig. 1d**). All unwinding activity ceased upon transfer to a solution lacking Ded1p and resumed upon returning to the solution containing Ded1p. This indicates that repeated binding and dissociation of either individual Ded1p protomers or complexes of Ded1p is required for repeated RNA hairpin unwinding and rezipping reactions.

**Figure 1:**
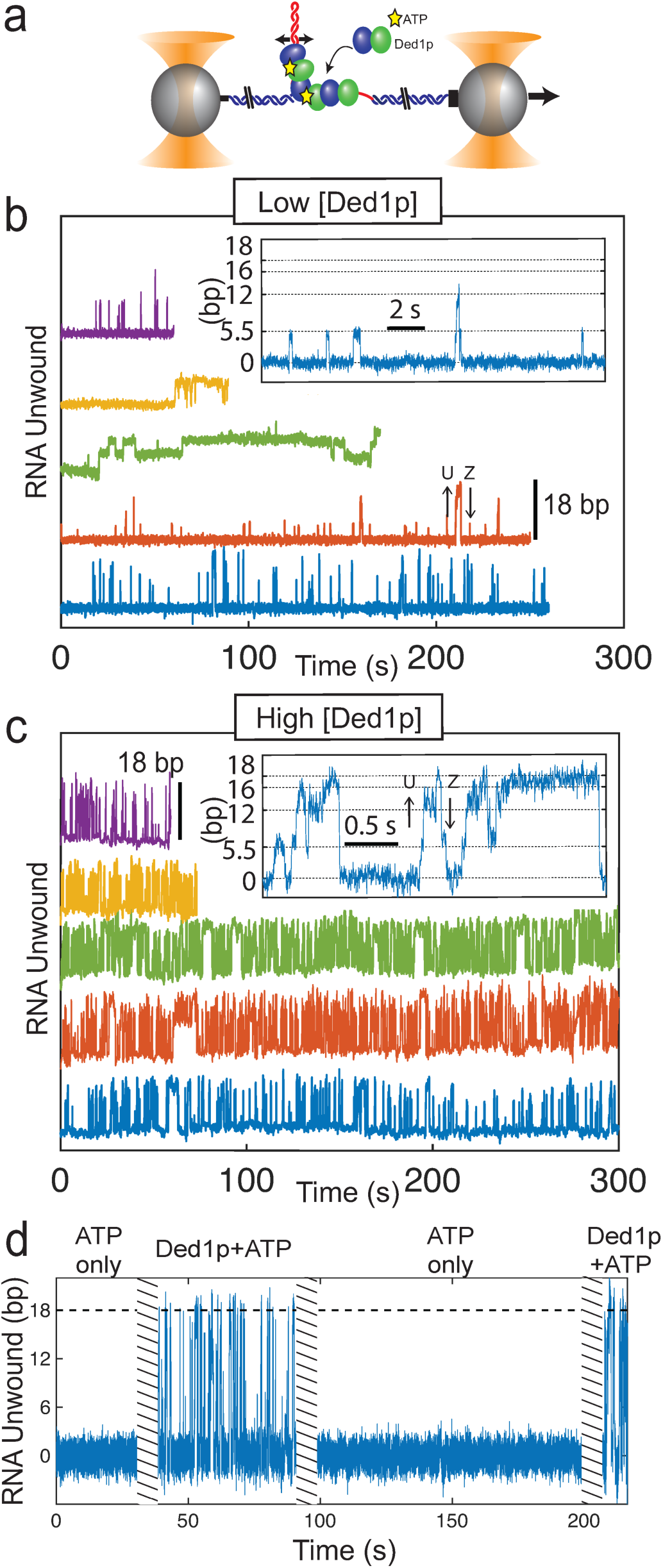
High-resolution optical tweezers measurements of Ded1p unwinding a 16 bp RNA hairpin with an adjacent 25 nt 3’ overhang. a) Schematic of dual-trap optical tweezers assay. ATP (yellow stars) and Ded1p protomers (blue and green oval balls representing Rec-A1 and A-2 domains respectively) bind and unwind an RNA hairpin with a tail (red) that is held on each side by 1.5 kb dsDNA handles, attached to a pair of polystyrene beads (gray spheres) each held at the focus of a laser (the optical traps; orange cones). The position of the trap on the left is fixed while the trap on the right moves left and right tracking the changes in extension of the RNA and maintaining a constant force of 10 pN for all experiments. b) and c) Five representative tweezers assay time trajectories each, offset vertically for clarity, showing RNA unwinding (increases compared to baseline; some example individual unwinding steps labeled U) and rezipping (decreases; some examples labeled Z) in the presence of 2.5 mM ATP and Ded1p at b) ‘low’ (100 nM) and c) ‘high’ (500 nM) concentration respectively. Insets show zoom-in examples with dashed lines to guide the eye corresponding to mean state locations as presented later. 18 bp corresponds to the maximum expected observation (16 bp stem and capping tetraloop). d) Tweezers assay time trajectory where a single RNA hairpin tether is transferred between a buffer containing only 2.5 mM ATP (i.e., no Ded1p; initial interval from t = 0 to t = 35 s, and third interval from t = 100 to t = 200 s) and an adjacent buffer containing both 2.5 mM ATP and high concentration 500 nM Ded1p (second interval, t = 45 to t = 95 s etc). Hatched bars cover the tether transfer between solutions where accurate tweezers measurement is not possible. 1.5 ms per data point for all plots.

We were able to build a detailed state model of the stepwise unwinding of RNA by Ded1p due to the high resolution of the tweezers combined with the high repetition rate of the reaction yielding very high statistics (individual tethers yielded between hundreds and thousands of repeated full unwinding cycles). For each unwinding reaction, we first found the unwinding and rezipping steps (changepoint-based step finding analysis; see Methods for details on all analysis) that thus defined the intervening RNA -Ded1p complex states (two examples for high Ded1p concentration reactions with their state model are shown in **Fig. 2a**; all the analysis results in **Fig. 2** are representative of high concentration Ded1p experiments, and are shown for a single example tether measured for a duration of 1750 s, yielding approximately 1800 unique unwinding reactions, with the exception of **Fig. 2e** which represents the combined results from multiple tether replicates). The unwinding step size distribution (**Fig. 2b** where step sizes >0) ranged from +2 to +12 bp, with two prominent peaks observed at approximately +5 to +6 bp (around two-third of steps population) and +2 to +3 bp (approximately one-third of steps population). Notably, the larger unwinding steps of +5 to +6 bp occurred near the beginning of the reaction while the smaller steps of +2 to +3 bp occurred near the completion of unwinding (**Fig. 2a & c-e**). The full RNA hairpin was very rarely observed to unwind in a single step. In contrast, the rezipping step size distribution (**Fig. 2b** where step sizes <0) ranged over the full apparent length of the RNA hairpin from −2 to −18 bp, but also with the same two prominent peaks at approximately −5 to −6 bp and −2 to −3 bp. Complete rezipping in a single step could occur from any partially unwound state.

**Figure 2:**
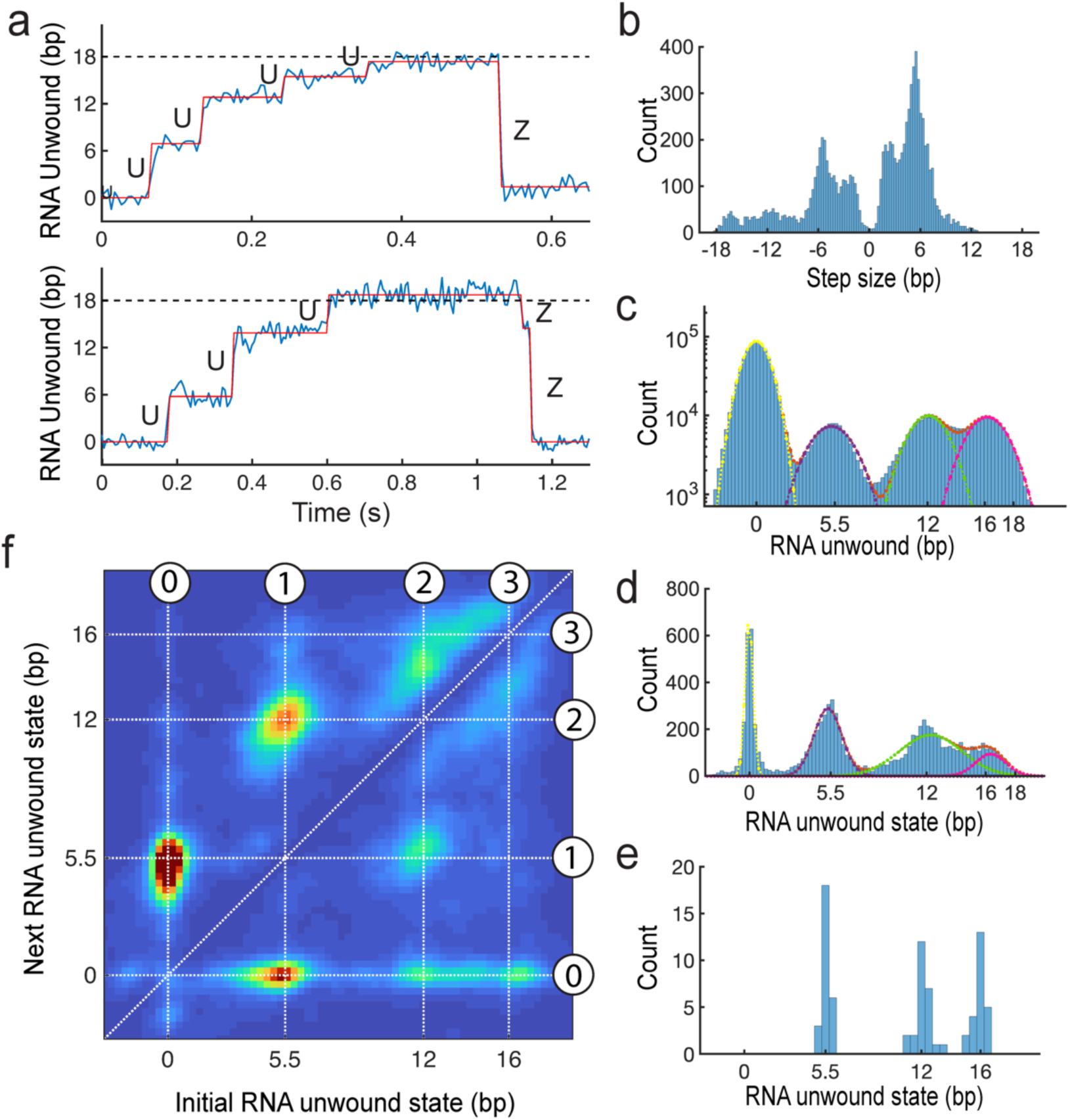
Detailed identification and analysis of state progression of Ded1p unwinding and rezipping of RNA at high Ded1p concentration (300 nM) and 2.5 mM ATP. All panels are derived from analysis of a single representative tether measured for 1750 s and containing 1800 unique unwinding reactions (i.e., two examples are shown in (a)), except for e) which combines results from multiple tether replicates. a) Two representative tweezers assay time trajectories zoomed in around single complete unwinding reactions showing raw data (blue, 1.5 ms per point) and the denoised states resulting from the changepoint-based state finding analysis (red). U and Z label individual unwinding vs rezipping steps respectively. Dashed line to guide the eye indicates the expected fully unwound hairpin distance. b) Unwinding and rezipping step size distributions (i.e., the distribution of differences between successive states). c) Distribution of raw tweezers assay data (e.g., the blue data shown e.g., in a)) with fit to a sum of four Gaussians (orange). Individual Gaussians making up the sum are also shown. d) Distribution of states of RNA unwinding by Ded1p found by analysis (e.g., the red fits shown e.g., in a)) with fit to a sum of four Gaussians (orange). Individual Gaussians making up the sum are also shown. e) Distribution of mean state positions (i.e., the location of the individual Gaussian fits to the state distributions in d)) for n = 25 unique tethers and replicated experiments. f) Distribution of Ded1p unwinding and rezipping state transitions. Heat map color shows the relative populations of each transition with red highest and blue lowest. The major states are labelled from 0 to 3 progressing from fully zipped to fully unwound with corresponding dashed lines to guide the eye. Transition density to the upper left of the diagonal guide are unwinding steps while density to the lower right are rezipping steps.

The unwinding steps correspond to a clear progression of states. **Figure 2c-e** show the distribution of states with increasing analysis precision. **Figure 2c** shows the distribution of raw tweezers data. Four major peaks are clearly observed, well-fit by a sum of four Gaussian distributions with the same widths as would be expected for a tweezers measurement of four major states (widths are approximately ±1.5 bp, for 1.5 ms integration time at 10 pN applied force, corresponding simply to the expected Brownian motion of the trapped beads). The first peak centered at 0 bp corresponds to the fully zipped hairpin (the initial state), while the following three states at approximately 5.5 bp, 12 bp, and 16 bp represent progressive unwinding of the hairpin (note that the standard error for each state position is negligible given the ±1.5 bp standard deviations and n greater than 100,000 data points per state distribution). **Figure 2d** shows the state distribution derived from the detailed step analysis. The 0 and 5.5 bp states representing the initial state and first partially unwound state are quite distinct and sharply defined compared to the following state distributions peaked at 12 and 16 bp which blend into each other. Hence this ‘denoised’ state analysis shows that there is a broader distribution of states near the completion of unwinding and not merely three states (as the noisy raw data might suggest). **Figure 2e** shows the distribution of state locations, when approximating by fitting with three Gaussians, over all unique tethers (i.e., experiment replicates; **Fig. 2d** is an example of a single tether). The means of these three well-separated peaks are 5.5±0.3, 12±0.5 and 16±0.4 bp respectively and are the best estimates of the dominant unwinding states (the standard error of the experiment replicates is given, which best represents the full experimental uncertainty). The unwinding states are not equally spaced in agreement with the heterogeneous step size distribution (**Fig. 2b**) and consistent with the non-translocating helicase model. The 12 bp state is in fact also well isolated and distinct from the approximately 16 bp state cluster. This can be seen in the state transition distribution plot in **Fig. 2f**, where each pair of successive states in the state progression are plotted as the next state vs the initial state. Here we see clear isolated peaks for both the first (labeled state 1 at 5.5 bp) and second (labeled state 2 at 12 bp) unwinding states. Unwinding always begins with a step from 0 bp to 5.5 bp. Next, unwinding either proceeds with a second step forward to 12 bp or reverses back to 0 bp. From the 12 bp state, two rezipping transitions are possible, either a reversal back to 5.5 bp or a direct transition to full rezipped at 0 bp. However, from the 12 bp state forward unwinding steps proceed differently compared to the first two unwinding steps: there is a final collection of states spaced approximately 1 bp and ranging 14 to 18 bp. Approximately 2 to 3 bp unwinding or rezipping transitions can occur between them. While a single step transition back to fully zipped (0 bp) can occur interestingly rezipping to the 5.5 bp intermediate is not observed.

Since the tweezers assay observed three progressive states of RNA hairpin unwinding, and previous ensemble experiments suggested a Ded1p trimer performs unwinding, it is tempting to interpret the three-step unwinding progression as the simple successive unwinding activity of three Ded1p protomers. However, note that the first two unwinding steps are distinct and highly reversible whereas the final apparent step is a collection of small transitions. Further, these final small transitions cover the apparent final 4 bp of hairpin opening, while it is unlikely that 4 bp or less of hairpin is stable on its own and available to be unwound (i.e., that amount of hairpin would spontaneously unwind). It is likely that the first two unwinding steps correspond to bonified unwinding of the hairpin by Ded1p, whereas the final subsequent smaller steps may correspond to some relaxation of the Ded1p and hairpin complex. The following detailed analysis of the Ded1p and ATP concentration dependent rate constants connecting the states supports these conclusions.

### Kinetic analysis suggests binding of successive single Ded1p protomers leads to successive steps

We performed tweezers measurements and detailed unwinding state analysis for varying Dedp1 concentration from below to above the ensemble suggested trimerization transition to understand how Ded1p binding to the RNA duplex corresponds to unwinding activity. For all concentrations, we observed the same unwinding states 0 through 3 (as labeled in **Fig. 2f**). However, at low Ded1p concentration (100 and 200 nM) the occurrence of states 2 and 3 was rare, consistent with the rezipping rate constant for state 1 → state 0 being nearly 10-fold larger than the unwinding rate constant for state 1 → state 2. At high concentration (300 and 500 nM) the state 1 → state 2 unwinding rate constant was the same or higher respectively than the state 1 → state 0 rezipping rate constant, consistent with the common observation of complete unwinding. Overall, we observed that rate constants for both the first two unwinding steps, i.e., state 0 → state 1 and state 1→ state 2, increased with increasing Ded1p concentration from 100 nM to 500 nM (**Fig. 3a** and **Supplementary Fig. 2)**. The increase in each individual unwinding step rate constant with increasing Ded1p concentration is consistent with individual Ded1p protomers binding with each unwinding step. If all the necessary protein (e.g., a full trimer) were already bound at the RNA hairpin when the first unwinding step occurred, the second step should show no Ded1p concentration dependence. The rate constants for all the rezipping transitions (1®0, 2®0, 2®1 and 3®0) decreased with increasing Ded1p concentration consistent with additional protomer binding suppressing rezipping. Notably, the final unwinding step (state 2 → state 3) follows the opposite trend as the first two unwinding steps and instead matches the concentration dependence associated with protomer dissociation.

**Figure 3:**
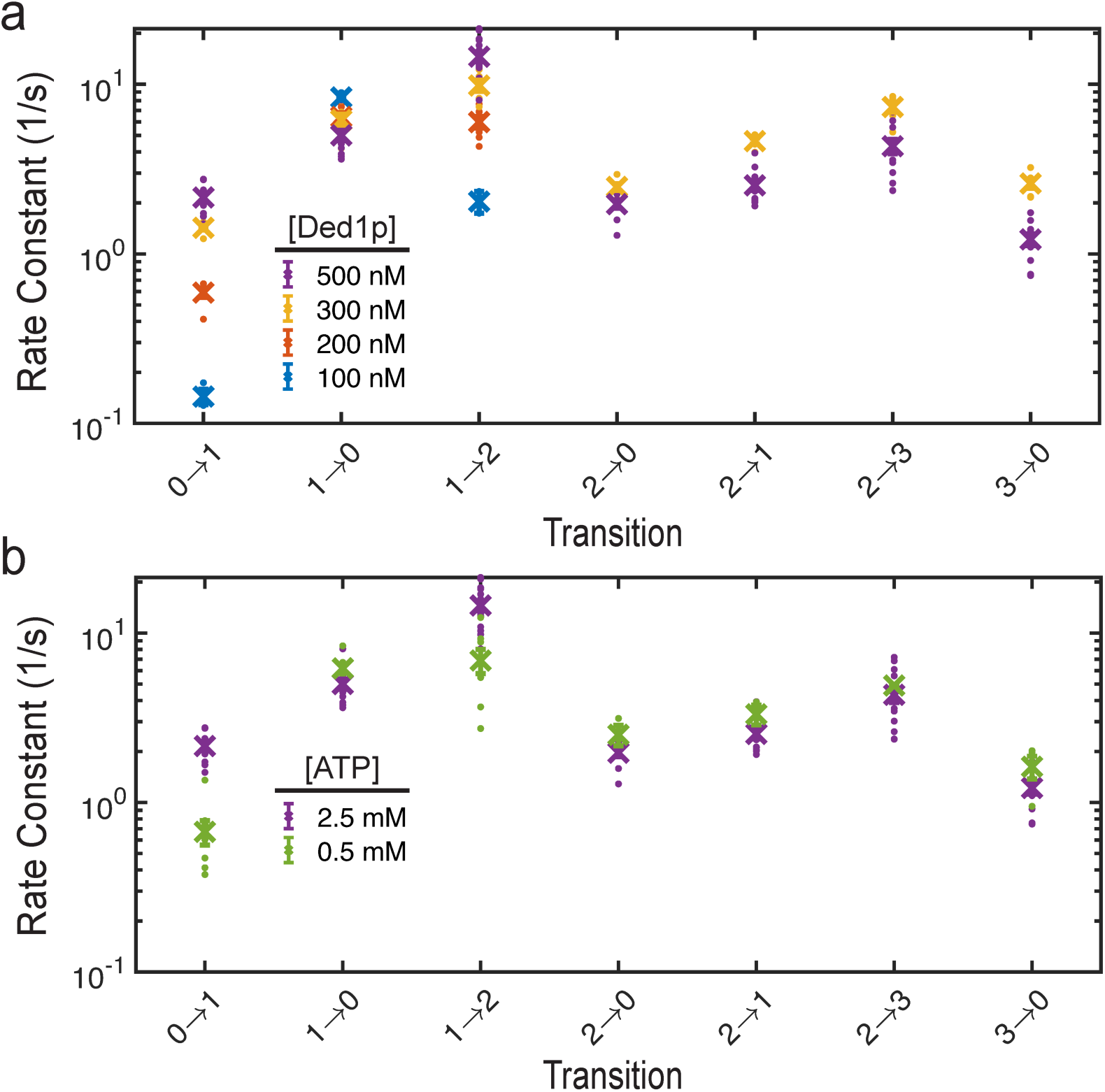
Ded1 unwinding and rezipping rate constants derived from detailed state analysis and their variation with a) Ded1p and b) ATP concentrations respectively. The unwinding transitions, as labeled on the x-axes, are indicated by increasing state number and correspond to the states labeled in Fig. 2 (see also Methods) as 0Ò1 (0Ò5.5 bp), 1Ò2 (5.5Ò12 bp), 2Ò3 (12Ò16 bp). Rezipping transitions correspond to decreasing state number. The symbol ‘x’ indicates the best estimate of the rate constant from all dwell times combined from multiple unique tethers, with error bars (often smaller than the ‘x’) corresponding to the standard error of the mean. The dots spread about the ‘x’ are rate constants derived from individual tethers (of unequal measurement durations) and give an alternate (overestimate) of measurement variance. A) The ATP concentration is fixed at 2.5 mM while tweezers assay experiments were performed at 100 (blue), 200 (red), 300 (yellow) and 500 (purple) nM Ded1p concentration. At the lower 100 and 200 nM Ded1p concentrations, insufficient transitions to state 2 occurred to be able to determine rate constants related to leaving state 2, or state 3, and thus they are not shown. B) The Ded1p concentration is fixed at 500 nM while tweezers assay experiments were performed at 0.5 (green) and 2.5 (purple) mM ATP concentration. Note that the data corresponding to 2.5 mM ATP is identical to the corresponding data in a).

We investigated the role of ATP in the stepwise unwinding of RNA via tweezers measurements and performed detailed unwinding state analysis for varying ATP concentration (0.5 and 2.5 mM) at high Ded1p concentration (500 nM; **Fig. 3b**). We observed an increase in the rate constants of the first two unwinding steps (0®1 and 1®2) with increased ATP concentration consistent with the necessity of a bound ATP for unwinding. The rate constants of the rezipping transitions (1®0, 2®0, 2®1 and 3®0) all decreased slightly with increased ATP concentration. Rezipping should occur after dissociation of Ded1p protomers from the RNA hairpin. The slight suppression of rezipping with increased ATP is consistent with ATP hydrolysis preceding dissociation and suggesting that ATP from solution can swap with ADP for Ded1p-ADP still bound to the RNA, thus re-stabilizing the helicase bound to the RNA. Note that again the final putative unwinding step from 2→3 behaves more like a protomer dissociation step, with the rate constant decreasing with increasing ATP concentration. We also performed tweezers experiments with the slowly hydrolysable ATP-analog AMPPNP instead of ATP. We observed similar multiple-step unwinding, however rezipping steps were strongly suppressed (**Supplementary Fig. 3**).

Overall, these kinetic results support the simple model that two individual Ded1p monomers, each with a bound ATP, bind and unwind the RNA hairpin sequentially, one after the other (corresponding to transitions from state 0 → state 1, then state 1 → state 2). What at first glance might also appear to be a final small unwinding step (state 2 → state 3) is likely not. The final step is likely a partial relaxation of the unwound RNA upon partial unbinding of Ded1p. However, ensemble experiments strongly suggested that three Ded1p proteins were necessary for efficient full unwinding and that the adjacent 3’ ssRNA tail was critical. We hypothesized that a third protein is bound to the tail and performed the following combined fluorescence and tweezers experiments to confirm.

### Fluorescence reveals a single Ded1p binds to the RNA tail to promote unwinding

While Ded1p binding and unwinding of the tethered RNA hairpin results in a tether extension increase and a readily detectable signal in the high-resolution tweezers assay, binding to the adjacent 3’ ssRNA tail did not result in a readily detectable tweezers signal. Therefore, to test for Ded1p binding to the tail we devised a single molecule fluorescence assay based on the protein-induced fluorescence enhancement (PIFE) method^47,48^ combined simultaneously with the tweezers unwinding measurements. We attached a Cy3 fluorophore to a base located in the middle of the 3’ tail (**Fig. 4a** and Methods for details). We expected to see an increase in Cy3 fluorescence intensity if a Ded1p protein binds nearby on the tail. **Figure 4b and Supplementary Figure 4** show example results: in buffer containing Ded1p (500 nM, i.e., high concentration) and ATP (2.5 mM), tweezers measurements showed repeated unwinding and rezipping of the RNA hairpin (same as above), while simultaneous fluorescence measurements showed intervals of increased fluorescence signal (approximately 1.5-fold increase, similar to previous PIFE applications^47^) strongly correlated with intervals of RNA hairpin unwinding activity. The fluorescence vs time signal was fitted by a two-state Hidden Markov model (HMM) and the tweezers data was analyzed to determine the procession of unwound RNA states as above. In **Fig. 4c** we plot the distributions of RNA unwound states (tweezers data) separated by whether the fluorescence state model was high or low, i.e., with or without a Ded1p bound to the tail. We observed that the low fluorescence state, indicating no Ded1p on the tail, occurred mostly when the RNA hairpin was fully zipped (state 0) and for this state high fluorescence was observed only 18±2% of the time (standard error via bootstrapping, as for all following errors on high fluorescence occurrence percentages). In contrast for the first two unwound states 1 and 2, fluorescence was majority high, with a Ded1p on the tail 72±3% and 63±3% of the time spent in states 1 and 2, respectively (**Fig. 4d**). **Figure 4e** shows the fraction of events with high fluorescence before and after the initial unwinding step (i.e., state 0 to state 1) as a function of time. Well before the unwinding step the fraction remains steady at ∼20% (i.e., approximately the overall average for state 0). However, at the moment of unwinding about 60% of the time fluorescence was high i.e., a protein was bound to the tail. After the unwinding step the fraction continues to grow to nearly 90%. Thus, while a protein bound to the tail enhances the probability of a protein binding to the hairpin and initiating unwinding, conversely the presence of the first protein bound to the hairpin also enhances the probability of a protein binding to the tail. Ultimately, the first step of unwinding consists of two proteins, one on the tail and one on the duplex.

**Figure 4:**
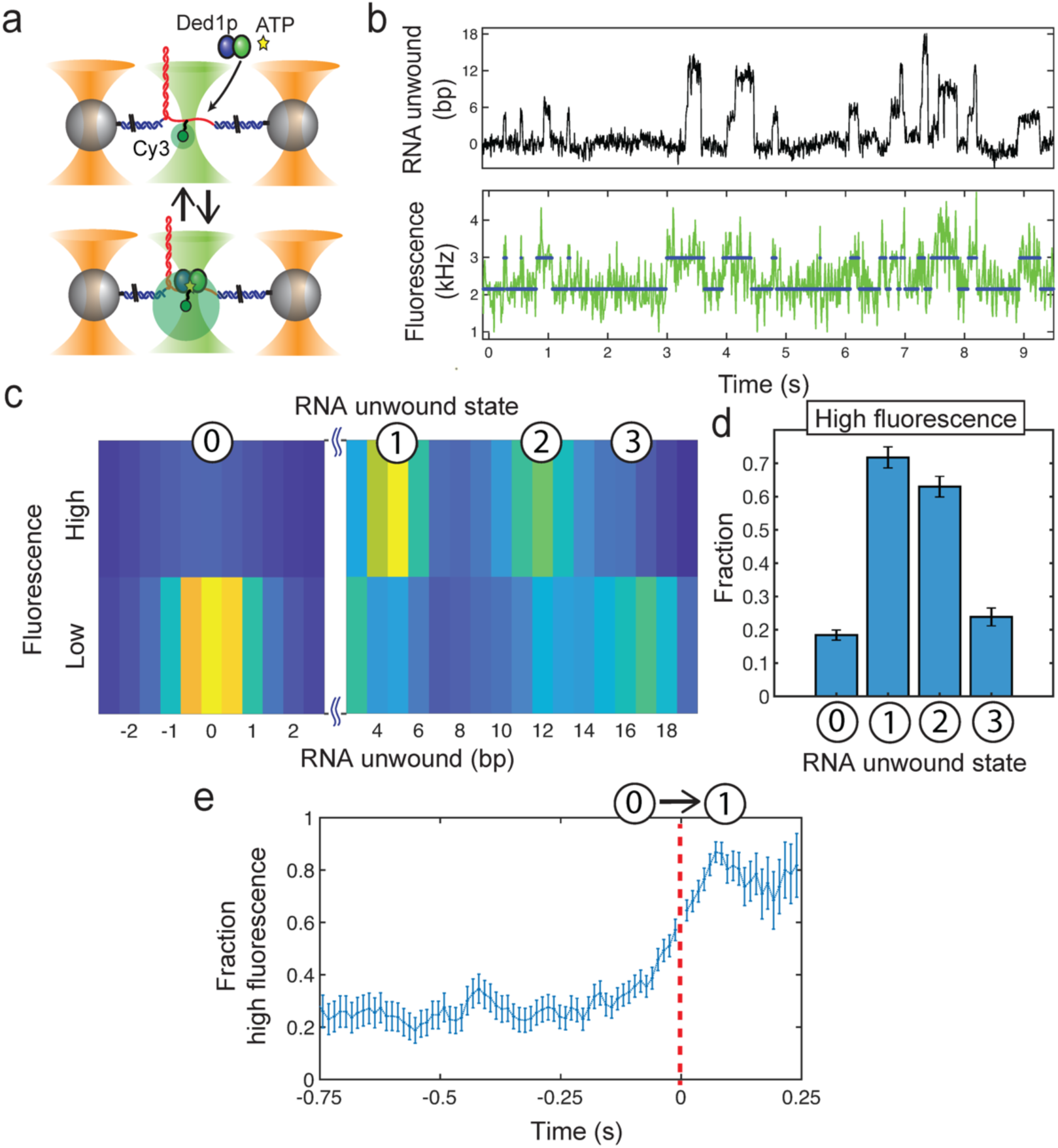
Assay combining high-resolution tweezers and single molecule confocal spectroscopy to measure Ded1p unwinding of hairpin RNA and Ded1p binding to adjacent ssRNA tail simultaneously. All data and analysis correspond to high Ded1p concentration (500 nM) and 2.5 mM ATP conditions (n = 34 unique tethers). a) Schematic of combined assay with tweezers assay as in Fig. 1a but with a single Cy3 fluorophore covalently attached to a base in the middle of the adjacent tail (green disks) and confocal laser excitation (green cones) and detection centered on the tethered RNA substrate to measure a PIFE signal upon Ded1 binding the tail. b) Example time trajectory with upper: standard tweezers unwinding assay as shown in e.g., Figs. 1 and 2, and lower: simultaneous PIFE fluorescence measurement (green) with 2-state HMM analysis results superimposed (blue). c) Distribution of RNA unwound state for intervals when fluorescence (i.e., PIFE) was high (upper) vs low (lower). Yellow vs blue indicates high vs low relative occurrence. State labels 0 to 3 are as previously defined (e.g., see Fig. 2). d) Fraction of each state population corresponding to high fluorescence (e.g., fluorescence was high for state-0 approximately 20% of the time). Error bars represent the standard error of the mean. e) Time correlation analysis of the specific transition from state-0 to state-1 (i.e., the first unwinding step) showing the fraction of events with high fluorescence as a function of time before and after the step occurrence. All events are aligned in time such that t = 0 corresponds to the time the unwinding step occurred (red dashed line to guide the eye; labels 0 and 1 indicate the states before vs after the step respectively).

However, the tail bound protein dissociates over time, leading to the slightly lower high fluorescence fraction for the final unwound state 2. The population of unwound states with high fluorescence decreased sequentially from 72±3% to 63±3% to 24±3% for states 1, 2, and 3, respectively (**Fig. 4d**). We rejected the null hypotheses that the state 1 and state 2 fluorescence distributions were identical and similarly that the states 0 and 3 fluorescence distributions were identical with greater than 97% and 96% confidence respectively using both bootstrapping methods and 2-sided T-tests which agreed. Unlike states 1 and 2, state 3 is predominantly associated with low fluorescence (**Fig. 4c and d**) indicating the absence of Ded1p on the tail. This is aligned with the rate constant vs protein concentration analysis above which showed that transitions to state 3 were consistent with protein dissociation. Again, state 3 consists of a set of configurations approaching the expected nearly fully unwound RNA extension. It’s important to note that state 3 can’t correspond to the RNA fully unwound with all Ded1p dissociated as the bare ssRNA would completely rezip to state 0 faster than our tweezers could observe (∼< 1 ms). State 3 likely comprises configurations where one or two of the Ded1p unwinding triplet have dissociated while the last Ded1p protomer to bind and unwind, near the RNA hairpin stem loop, is still bound and preventing the RNA hairpin from rezipping (**Supplementary Fig. 5)**. Unbinding of Ded1p should occur after ATP hydrolysis. Since ATP hydrolysis is stimulated by being bound to the RNA, the Ded1p protomers that bind first will also tend to dissociate first. This is consistent with the tail bound Ded1p dissociating first leading to the low fluorescence signal dominating state 3.

## Discussion

In this report, we have explored the mechanistic details of cooperative RNA unwinding by Ded1p helicases at the single-molecule level using a high-resolution fleezers assay. We observed a dramatic increase in unwinding activity at high Ded1p concentration, consistent with previous ensemble experiments^19^. At high Ded1p concentration, the RNA substrate underwent thousands of repeated cycles of unwinding and rezipping that required the continued presence of both Ded1p protein and ATP in solution, indicating the requirement of binding and dissociation of proteins for each cycle. This contrasts with an alternate model where a single complex of Ded1p assembles on the RNA which is capable of repeated full unwinding and rezipping without dissociation. Our tweezers measurements resolved the time progression of four major states, states 0 – 3, and the rate constants of transitions between states as a function of both Ded1p and ATP concentration. Simultaneous fluorescence measurements revealed binding of Ded1p specifically to the 3’ ssRNA tail adjacent to the RNA duplex. State 0 consists of the RNA fully zipped with a Ded1p sometimes bound to the tail (**Fig. 5**). State 1 occurs after the first step of RNA unwinding consistent with a single Ded1p protomer binding presumably to the stem region of the RNA duplex. That protein binding is concurrent with unwinding is indicated by the nearly linear increase in the first step unwinding rate constant with increasing Ded1p concentration. Usually, the first unwinding step is preceded by binding of a Ded1p to the adjacent tail or is quickly followed by binding of a Ded1p to the tail, indicating that two Ded1p are bound to the duplex and tail together in state 1. Rezipping from state 1 back to state 0 occurs frequently and is independent of protein concentration, consistent with ATP hydrolysis, and indicates protein dissociation. Progression from state 1 to state 2 in the second unwinding step unwinds the remaining RNA duplex and is also consistent with the concurrent binding of a single Dedp1 protomer. As with the partially unwound state 1, a Ded1p protomer is also usually bound to the adjacent tail. Thus state 2 consists of the RNA duplex fully unwound with two Ded1p protomers bound to the duplex and one bound to the adjacent tail. State 2 frequently reverses back to state 1, consistent with the individual dissociation of the third Ded1p protomer. State 3 is a collection of configurations near the apparent extension of the fully unwound and bare RNA duplex that are consistent with the dissociation of either the first (i.e., tail bound) or second (i.e., duplex stem bound) Ded1p individually and the resulting reverting of those portions of substrate to bare RNA. Thus, complete unwinding of the 16 bp long RNA duplex is a highly dynamic and stochastic process of three Ded1p protomers individually binding and dissociating directly on the RNA duplex and adjacent tail substrate.

**Figure 5:**
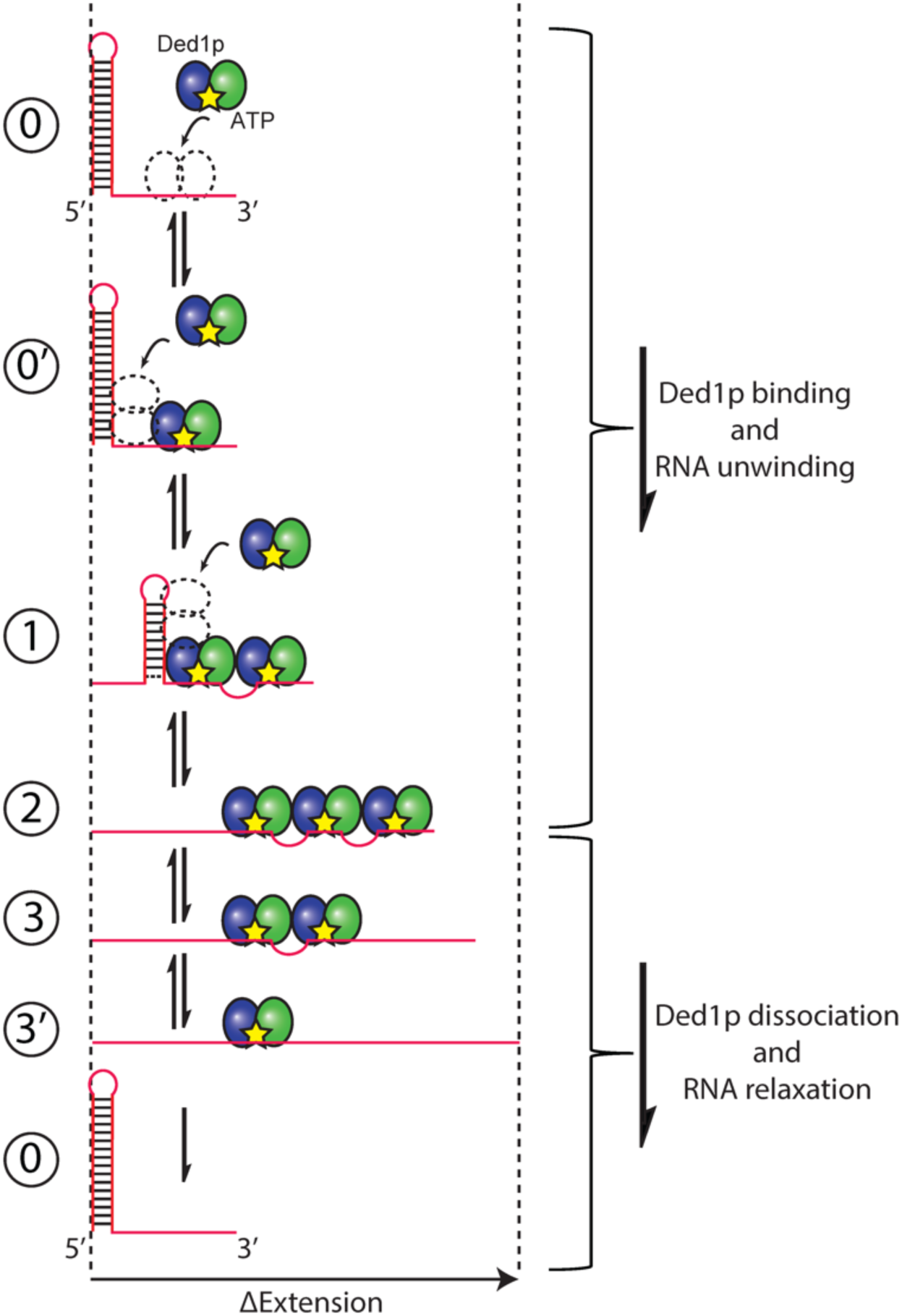
Mechanistic model of the major pathway of RNA unwinding by a set of three Ded1p helicases. Individual Ded1p protomers illustrated with Rec-A1 and Rec-A2 domains as blue and red ovals respectively and ATP as yellow stars. The reaction initiates at the top of the diagram and proceeds downwards. Major states are numbered on the left and correspond to the previous descriptions in the text. The extension of the RNA substrate as would be detected by the tweezers measurements is portrayed horizontally. State 0 and 0’: RNA hairpin begins fully wound and with no bound Ded1p (state 0) and then a single Ded1p binds to the adjacent 3’ ssRNA tail (state 0’). No change is extension is observed though state 0’ is distinguished from state 0 via measurement of high fluorescence (high PIFE). State 1: a second Ded1p has bound and unwound a portion of RNA hairpin. The first and second Ded1p may interact, leading to a small contraction of the unwound RNA as depicted by the small ssRNA bubble. State 2: a third Ded1p has bound and unwound the remaining RNA hairpin. The third Ded1p may interact with the second leading to a small contraction of the unwound RNA. State 3: the first Ded1p has dissociated from the tail, releasing a small amount of contracted RNA, and increasing the extension. State 3’: the second Ded1p has dissociated, releasing a small amount of contracted RNA, and increasing the extension. After the third Ded1p dissociates the tether will fully rezip and return to state 0.

The amount of RNA unwound by the first Ded1p binding to the duplex in the first unwinding step, about 5 to 6 bp, is remarkably consistent (as shown in **Figs. 2d** and **e** state 1 distributions). **Fig. 2d** shows the distribution of state 1 extensions corresponding to a mean of 5.5 bp and a standard deviation of 1 bp. The variance is not simply due to the noise from the Brownian motion of the trapped beads (i.e., as seen in the raw data extension distributions in **Fig. 2c**), but actual variation in the amount of RNA that is unwound in the first step of unwinding. Note also that this measurement employs the standard hairpin assay interpretation of changes in tether extension, where the unwinding of a single base pair increases the tether length by two nucleotides. More accurately, the step size determination should also include additional contributions to the change in extension including deviations from the ideal ssRNA extension with Ded1p protein bound to it (e.g., possibly ‘scrunching’ of the ssRNA backbone with bound Ded1p) and/or interactions between adjacent Ded1p proteins (e.g., the first unwinding Ded1p protein interacting with a Ded1p on the adjacent tail; likely also leading to an additional contraction). Any scrunching of the ssRNA backbone is likely 1 bp or less and this effect would lead to a constant offset to the 5.5 bp measurement independent of the specific duplex binding location. Alternatively, Ded1p could bind ssRNA at additional protein binding sites, similar to how the Hepatitis C Virus helicase ns3 was shown to release nascent unwound RNA strands asynchronously resulting in apparent half bp unwinding steps,^49^ though there is no indication that the smaller Ded1p protein has such activity. Most likely, multiple Ded1p on the RNA transiently interacting or binding to each other would likely contract a variable amount of ssRNA determined by the exact binding location of each Ded1p and the subsequent amount of ssRNA in between that is contracted. As noted in the results above, the final small increases in tether extension approaching the fully unwound tether length (18 bp is the maximum that could be observed, consisting of the full 16 bp hairpin unwinding plus the release of the 4-nt stem loop) which are part of state 3, are consistent with protein dissociation from the substrate. There is a range of step sizes observed within state 3, from 1 to 3 bp, which suggests a range of ssRNA contraction between interacting Ded1p from 2 to 6 nt. In the end, the 5.5 bp mean amount of RNA unwound by a single Ded1p may be underestimating by 1 to 2 bp. Further some or all of the variance in the observed unwinding amount could be from the variation in specific Ded1p binding locations and subsequent variation in the amount of ssRNA contracted in between. While the second unwinding step apparently unwinds 6.5 bp, going from 5.5 bp unwound to 12 bp unwound, most likely the duplex is fully unwound after the second step. The hairpin is only 16 bp long, and it is extremely unlikely that exactly 12 bp could be unwound leaving the final 4 bp plus 4 nt stem loop stably wound. Again, as discussed above, direct interactions of Ded1p with the ssRNA or between Ded1p reduce the extension of the unwound ssRNA. Thus, while it is clear that a second Ded1p needs to bind and unwind to fully unwind the RNA, the approximate 5.5 bp unwinding by the first Ded1p should be the best estimate of how much RNA a single Ded1p can unwind. Future experiments investigating the detailed interactions between Ded1p proteins and a purely ssRNA substrate will be necessary to improve our understanding.

## ACKNOWLEDGEMENTS

M.J.C. acknowledges support from NSF (MCB-1919439). E.J. acknowledges support from the NIH (GM118088).

## METHODS

### Ded1p expression and purification

Recombinant Ded1p was expressed and purified in *E. coli* as previously described^33^.

### Optical tweezers RNA unwinding assay

All optical tweezers measurements were carried out at 23 °C using a home-built high-resolution dual optical trap combined with single molecule confocal fluorescence spectroscopy instrument as previously described in detail^18,50,51^. In brief, an acousto optic modulator (AOM) rapidly deflects a single 1064 nm laser between two locations within a microfluidic sample chamber to generate two independently controlled timeshared optical traps. Each trap laser intensity, and thus the trap stiffness, was stabilized by a PID-based active feedback method. Trap stiffness of 0.25 pN/nm was typical. Each trap bead position was measured once per timesharing cycle (i.e., once per 15 µs) by an infrared-enhanced quadrant photodiode using the standard back focal plane interferometry method. Time trajectory measurements (e.g., **Fig. 1b**) were performed using the active force feedback method where a constant force, 10 pN – well below the spontaneous opening force of the RNA hairpin, approximately 16 pN - was maintained via keeping the difference between the two trapped bead positions within the traps constant by adjusting the location of one movable trap. The trap position was adjusted by the feedback 100-fold slower than the bead measurement rate, every 1.5 ms. Thus, the position of the movable trap is the data that reports on the changing length of the tether, and thus the unwinding vs rezipping of the tethered RNA hairpin vs time. This change in tether extension measured initially as a distance is then converted to the number of corresponding base pairs of hairpin unwound according to standard methods and as described in detail below. The two traps on-time durations are 5 µs each and were followed by a 5 µs fluorescence measurement interval where the trap lasers were off and a fluorescence excitation laser (532 nm for the experiments in this paper) and single photon fluorescence emission detection (by single photon counting avalanche photodiodes) were on, giving an overall timesharing and interlacing repetition rate of 67 kHz (15 µs period). Fluorescence data was then integrated to the final values as detailed in the associated figure captions (typically ∼10 ms; see analysis method details below). Overall data acquisition and control are performed by an FPGA-based PC DAQ card (National Instruments PCIe-7852R) programmed via LabVIEW (National Instruments). Offline data analysis is performed as described in detail below.

Optical trapping tether constructs contained a single RNA/DNA chimeric insert, annealed, and ligated to two flanking dsDNA linkers referred to as handles and were generated using the general methods as described previously^17^. DNA (handle primers) and RNA (hairpin substrates) oligomers were acquired from IDT except for the fluorophore-labeled RNA hairpin which was acquired from Dharmacon (now Horizon). The central RNA portion of the insert consisted of a 16 bp duplex capped with a 4-nt loop along with an adjacent 25 nt ssRNA overhang at the 3′-end (**Supplementary Fig. 1**). Both the 5’ and 3’ ends of the central RNA were extended with ssDNA complimentary to overhangs generated on the two handles specifically. Thus, the only RNA in the tether is the RNA hairpin and adjacent 3’ ssRNA tail. For PIFE experiments, the RNA substrate sequence remained unchanged, but a modification was made in middle of the overhang (13^th^ nucleotide from 3’-end) by labeling Cy3 to the base of nucleotide (Ded1p is known to bind to the RNA backbone).

Briefly, the two handles of ∼1.56 kb and ∼1.69 kb, termed as left and right handles (LH and RH), were initially generated by PCR amplification of plasmid pBR322 and Lambda phage DNA to give ∼ 2.18 kb and ∼2.21 kb DNA pieces respectively. The forward primers were modified with biotin (for LH) and with digoxigenin (for RH) so that the final construct would tether on one end specifically to streptavidin-coated beads vs anti-digoxigenin-coated beads on the other. The amplified pieces were purified using a PCR clean-up kit (Qiagen) and then restriction-digested to create overhangs for ligation to the RNA substrate: LH was digested using either Hind III or PspGI and RH with TspRI. The cut product was purified using 1% (w/v) agarose gel, where the band corresponding to the desired handle length (∼1.6 kb or 1.7 kb for LH and RH respectively) was excised and then the DNA extracted using a gel-extraction kit (Qiagen). The RNA hairpin substrate was then annealed and ligated between the LH and RH handles (see final construct configuration as in **Fig. 1a** and **Supplementary Fig. 1**) following two slightly different protocols that yielded the same ultimate results, either: method 1) 5x excess RNA insert oligo with respect to an LH amount was combined with LH and ligated using T4 DNA ligase, the product was purified in 1% agarose gel as above, then finally this LH-RNA complex was ligated to an excess of RH and gel purified. Method 2: a single ligation reaction was performed with a 1:1:1 molar ratio of all three tether components (LH, RH, and RNA insert), and then purified via the 1% agarose gel method. The concentration of resulting ∼3.3 kb trapping construct was determined based on A260 nm absorbance on a Nanodrop spectrometer. The ∼3.3 kb construct was attached to 800-nm-diameter streptavidin-coated polystyrene beads (Spherotech) with a 1-hour room temperature and stored in 1x phosphate-buffered saline (PBS) supplemented with 0.001 % Tween-20 at 4 °C. The stored tether-coated-beads were diluted 200x in T50 buffer (consisting specifically of 20 mM Tris-HCl pH 8.0 and 50 mM NaCl) and injected into the corresponding sample chamber channel for each tweezers measurement session. ATP and AMPPNP were purchased from Sigma Aldrich.

Experiments were carried out within homemade sample chambers generated by sandwiching and melting (to adhere) laser-cut parafilm between two cover glass producing a 4-channel microfluidic chamber^18,51^. The two outermost channels are designed to supply two distinct types of beads: anti-digoxigenin-coated and tether-streptavidin-coated. Two microcapillaries connect the two bead channels to a pair of adjacent central channels where optical trapping and experiments are performed. The two adjacent center channels contain distinct solutions that do not mix while there is constant laminar fluid flow. In general, one solution channel contains a “blank”, control buffer (40 mM Tris-HCl pH 8.0, 50 mM NaCl, 2.5 mM MgCl2, 2 mM DTT, 2.5 mM ATP, unless specified otherwise in the presence of 1% glucose oxidase/catalase oxygen scavenger system to prevent oxidative damage to samples and premature tether breaking) while the second solution channel is identical except for the addition of Ded1p.^51^ For experiments including fluorescence measurement, 4 mM Trolox was also added. To prevent protein adhesion onto chamber surfaces in order to maintain accurate and stable Ded1p concentrations during experiments, chamber cover glass surfaces were passivated with mPEG-SVA, 5000 (Laysan Bio) as described in reference ^51^. To ensure smooth fluid flow that minimally interferes with high-resolution tweezers measurements, the central sample channels are aligned vertically, thus with vertical flow direction with respect to the horizontal arrangement of traps and beads and subsequent measurement (i.e., flow induced noise is nominally orthogonal to the measurement direction) and high precision and low noise syringe pumps (Harvard Apparatus, PHD2000 and Nano pumps) were used to inject and control fluid flow in the chamber. The chamber was mounted on a precision motorized stage (Newport), facilitating trap positioning within, and transitioning between either flow channel. Transitioning between flow channels took 1 to 2 seconds. Experiments were initiated with the *in situ* formation and validation of individual tethers between a pair of trapped beads. First, we trapped a single anti-digoxigenin-coated bead (i.e., not coated in the RNA tether construct) injected into the Ded1p-containing sample buffer nearby the linking capillary. Next, we moved to the adjacent blank channel nearby the capillary injecting RNA-tether-streptavidin-coated beads to capture the second bead in trap 2. This order of trapping ensured that the RNA-tethers were not prematurely exposed to Ded1p. Tethers were formed by the standard fishing method (oscillate one trap’s distance from the other near and far until a linkage is detected via a rising force upon withdrawing), and then validated by performing a standard force vs. extension measurement and comparing to the expected single tether curve. The expected single tether curve is modeled as the pulling curve of the sum of the 3.2 kb long dsDNA handles, plus the inline 25 nt long ssRNA (the 3’ tail adjacent to the RNA hairpin insert) and either with or without the 36 nt ssRNA contribution of the unzipped RNA hairpin (16 bp duplex plus 4 nt stem loop) corresponding to modeling the pulling curve with the RNA hairpin either zipped or unzipped which should correspond to the measured data in either the lower or upper force regions respectively. Polymer extension vs force modeling was done in the standard way with dsDNA modeled by the Extensible Worm-like Chain model (persistence length = 53 nm, stretch modulus = 1200 pN, distance per base pair = 0.34 nm) and ssRNA modeled by the Extensible Freely-Jointed Chain model (persistence length = 0.75 nm, stretch modulus = 800 pN, and distance per nucleotide = 0.59 nm^18^). Further, validated tethers needed to display the characteristic RNA hairpin unzipping force for this sequence and buffer conditions at ∼16 pN (**Supplementary Figure 1b**). All non-conforming tethers were discarded. After validating a single RNA hairpin tether via the force vs extension measurement, the constant-force feedback method was established with a force of 10 pN. The tether was observed in the blank buffer (i.e., no Ded1p present) for no less than 30 s to confirm the stability of the tether and lack of spontaneous unwinding and rezipping-like activity. Fully validated single tethers were then moved to the adjacent sample solution channel containing Ded1p and RNA hairpin unwinding was monitored. Additionally, in the case of PIFE fluorescence measurements simultaneous with tweezers measurements, the fluorescence excitation laser was turned on for a short duration in the blank buffer to verify the presence of a single, stable Cy3 fluorophore on the RNA insert. The Ded1p and ATP concentrations varied in different experiments.

### Data analysis

Offline data analysis was performed using automated custom MATLAB code (MathWorks). The end-to-end change in tether extension measured during the constant-force tweezers RNA hairpin unwinding assay experiments (the primary measurements presented in this paper, as in **Fig. 1b**) was converted to the equivalent quantity of RNA duplex unwound via the standard method of assuming two nt of ssRNA are added to the tether extension for each bp unwound. The expected change in extension, at a given force, for adding two nt to the tether is determined by the polymer modeling methods given above. The resulting calibrated time trajectory data were boxcar averaged depending on the particular analysis task to a final rate of between 1.5 to 24 ms per point. In general, unwinding data could be acquired for very long times for individual tethers (> tens of minutes often, with thousands of individual unwinding and rezipping cycles repeated). To combine all the individual reactions over the long duration measurements we employed a drift correction method to the data as follows. Histograms of 1 to 10 s segments of data (interval duration depending on drift rates and overall reaction conditions, i.e., the specific Ded1p and ATP concentration of the experiment), were produced and intervals showing no unwinding activity were identified (i.e., only a single peak in the tether extension distribution that was also nearby the preceding single peak that had been identified as the fully zipped state). A spline curve was produced that best fit only the intervals of no unwinding. Next, that spline curve was used to level only the data segments between unwinding reactions. To be clear, no drift correction was applied directly to unwinding data, but rather only the intervening fully zipped reference intervals. In this way the accumulated data from long measurement durations for single tethers could be aligned without fear of any distortion of the unwinding data. Following drift correction, a very simple changepoint processing was applied (the very efficient MATLAB provided algorithm) to identify steps in the time trajectory (the distribution of steps for one example tether is shown in **Fig. 2b**). This method required a threshold level which we set slightly above the bare tether noise level (as can be measured from the control data). The sequence of steps then defines a sequence of states. Each state in the sequence is labeled with its mean extension and its dwell time. The sequence of state means can be seen in red in the two examples of analysis in **Fig. 2a**. The distribution of state means for the same example tether is shown in **Fig. 2d**. The transition density plot derived from the same example tether and shown in **Fig. 2f** is the distribution of all sequential pairs of states from the state sequence, plotted using the standard kernel density method where a topographic distribution plot is produced by summing 2D Gaussians at the location of each state transition initial-final (x-y) pair. Individual state extensions differ from the peak values both for simple statistical reasons (i.e., a shorter duration state will have a larger extension measurement uncertainty) and because the exact state extension may also vary.

For the determination of the rate constants for the transitions between major states 0 through 3 (as seen in **Fig. 2f**), individual states from the state train were grouped into collections 0 through 3 according to being approximately within one sigma of the peak state location (as determined by the 4-Gaussian fitting shown e.g., in **Fig. 2c**). Hence, we excluded individual states whose extensions lie ambiguously between major states. Next, the mean dwell times of each major state were determined (along with the standard error of the mean) and then the rate constant for leaving that major state is simply the inverse (with the associated relative error). All dwell time distributions were consistent with single exponential distributions. We then determined the relative occurrence of transitions to specific final major states (i.e., reaction branching ratios), and combined with the overall rate constant for leaving the state (which is the sum of the rate constants for each branch), we thereby determined the specific rate constants to transition between the major states and their propagated relative error as plotted in **Fig. 3**.

For PIFE fluorescence experiments we adjusted the boxcar downsampling of the tweezers extension data and the integration time of the fluorescence data to give matching final data rates. We performed extension step and state analysis as above but in addition we performed standard 2-state HMM modeling on the fluorescence data. We then used the resulting extension and fluorescence state train models to produce the correlation and distribution plots described in detail in **Fig. 4**.

## Author Contributions

E.M.P., R.Y., M.J.C., and E.J. designed the experiments. E.M.P., R.Y., K.S. and K.C. performed the high-resolution optical trapping experiments. R.Y., E.M.P and M.J.C carried out the data analysis. A.A.P. expressed and purified Ded1p. R.Y., E.M.P., E.J., K.S. and M.J.C. wrote, reviewed, and edited the manuscript. All authors read and commented on the paper.

---

**Supplementary Figure 1:**
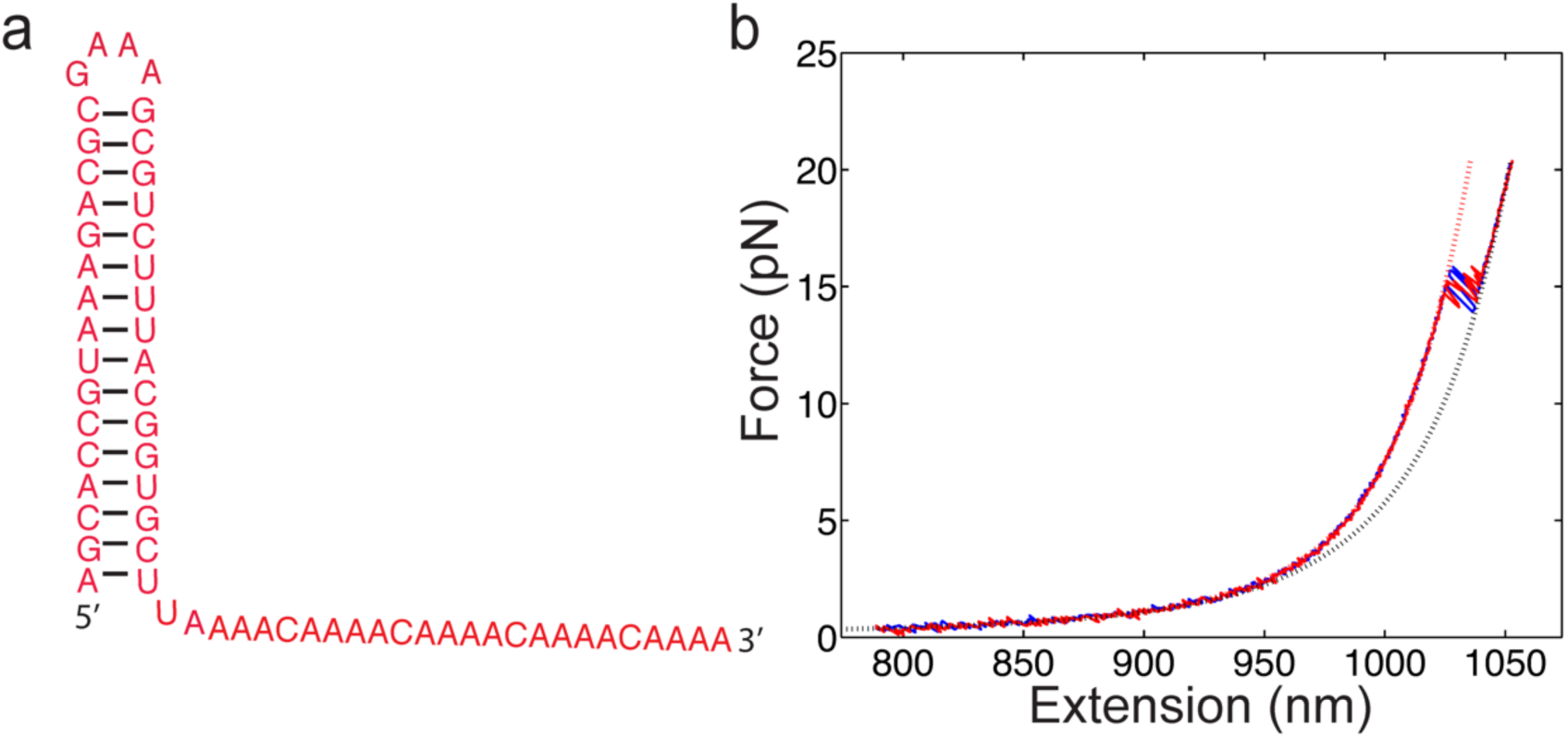
The RNA hairpin construct: A) The secondary structure of substrate RNA used in this investigation consisting of a 16 bp duplex RNA capped with a 4 nt (GAAA) loop and with an adjacent 25-nt 3¢ single-strand. The sequence of the substrate hairpin is the same as previously used in prior biochemical and biophysical studies^18^. The 4-nt cap serves to monitor the rezipping events by ensuring the duplex strands remain linked and are not lost after full unwinding. The ends of this RNA substrate were extended with DNA sequences complementary to the overhangs of the left and right tethering handles (LH and RH; see Methods for details). B) A measurement of the force vs extension of a representative good tether (blue solid line is a pull and red solid line is the following relaxation). Polymer models of the expected results for the RNA hairpin fully zipped (red dashed line) vs fully unzipped (black dashed line) shown by the dashed lines. The measurements overlap well with the models above and below the expected unzipping transition force, ∼15 pN.

**Supplementary Figure 2:**
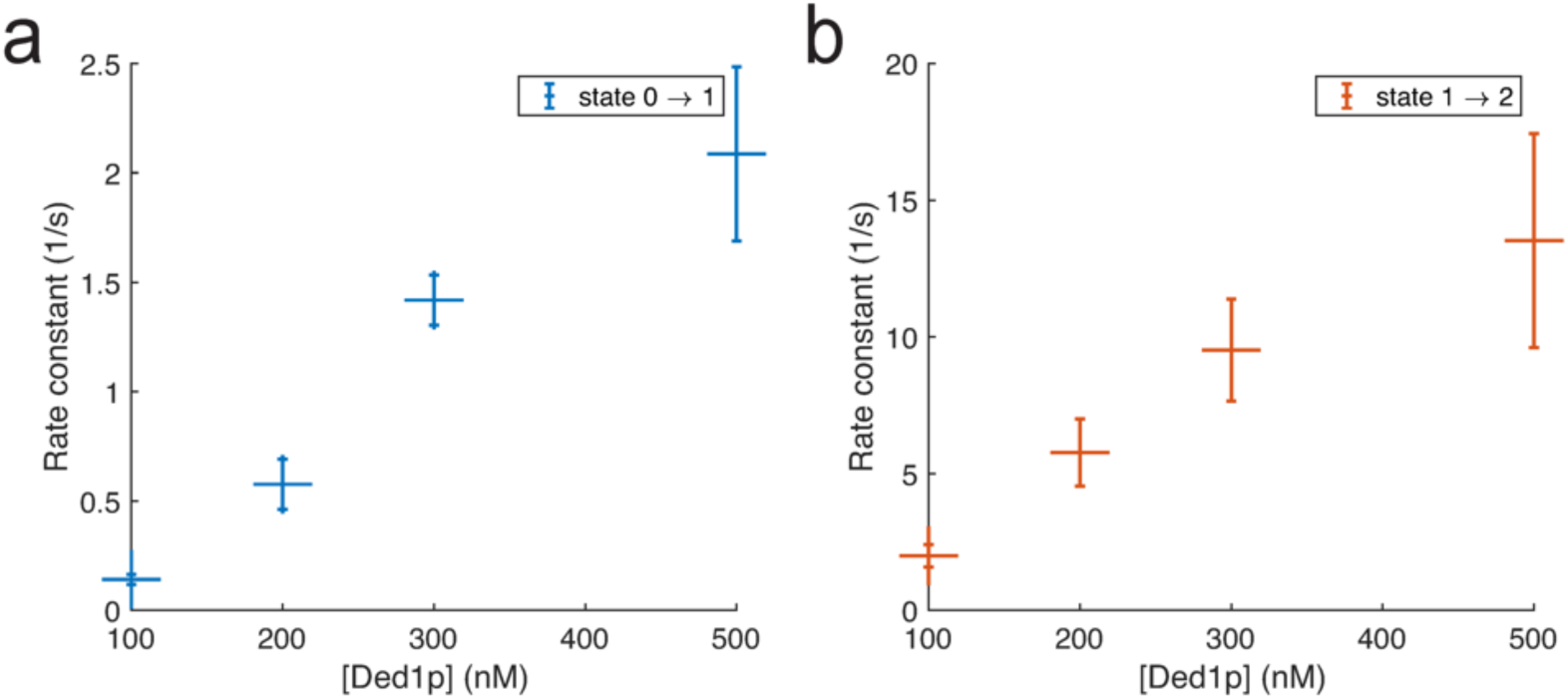
Zoom-in of Ded1 unwinding rate constants vs Ded1p concentration reproduced from Fig. 3a for a) the transition 0Ò1 (0Ò5.5 bp) and b) the transition 1Ò2 (5.5Ò12 bp). ‘+’ symbols indicate the best estimate of the rate constant from all dwell times combined from multiple unique tethers, with error bars corresponding to the standard error of the mean. The ATP concentration is 2.5 mM.

**Supplementary Figure 3:**
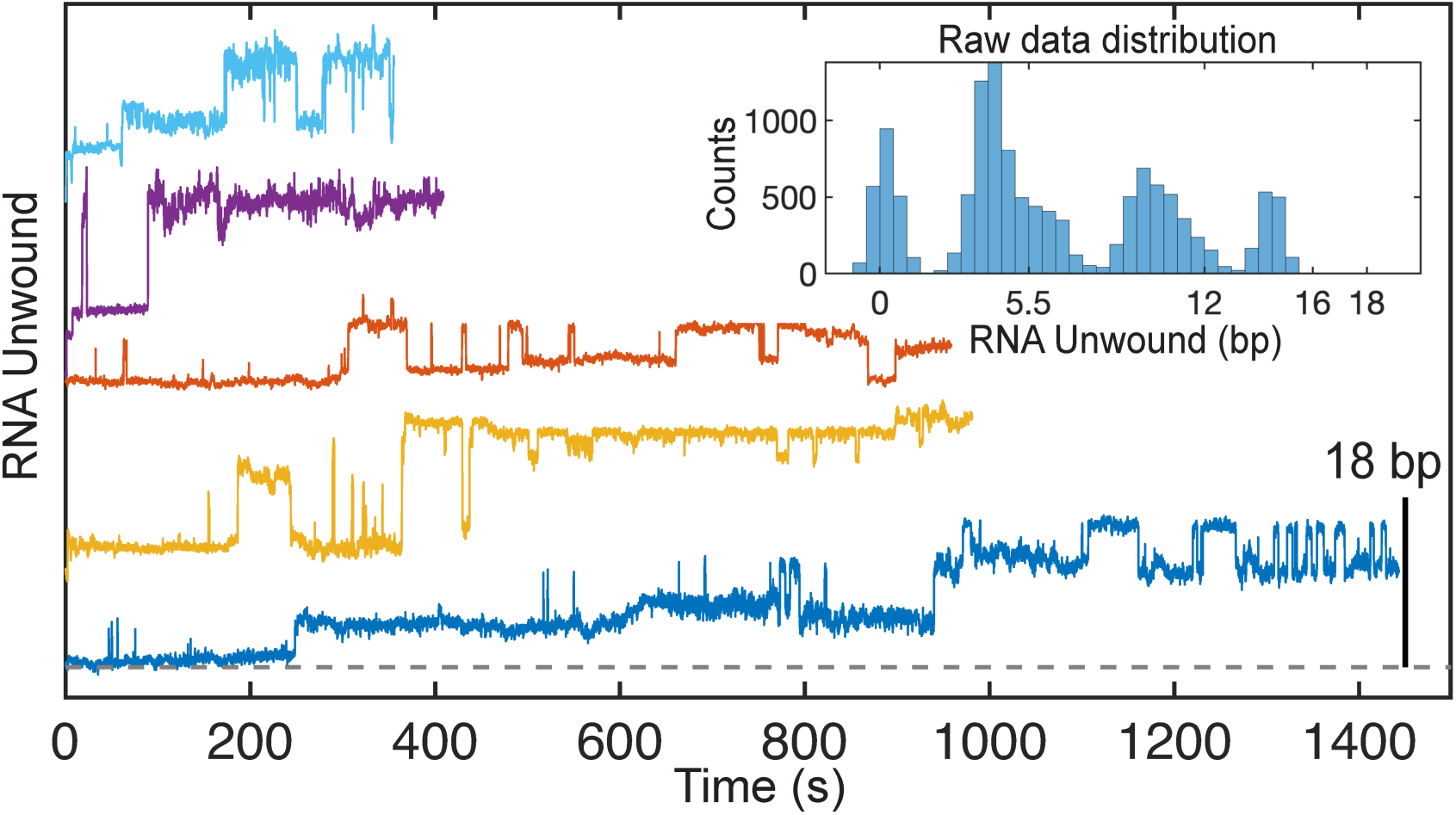
Representative tweezers measurements (vertically offset for clarity) of 300 mM Ded1p unwinding RNA in presence of 2.5 mM non-hydrolyzable nucleotide analog adenylyl-imidodiphosphate (AMPPNP). The maximum expected unwound extension corresponds to ∼18 bp as labeled by the vertical scale bar. Inset: Example distribution of raw unwinding data (generated from the bottom-most and longest time trace example in blue). Overall, while unzipping and unwinding activity is still observed in the presence of AMPPNP, the complete reaction time is much, much slower compared to standard conditions using ATP (as shown throughout the text).

**Supplementary Figure 4:**
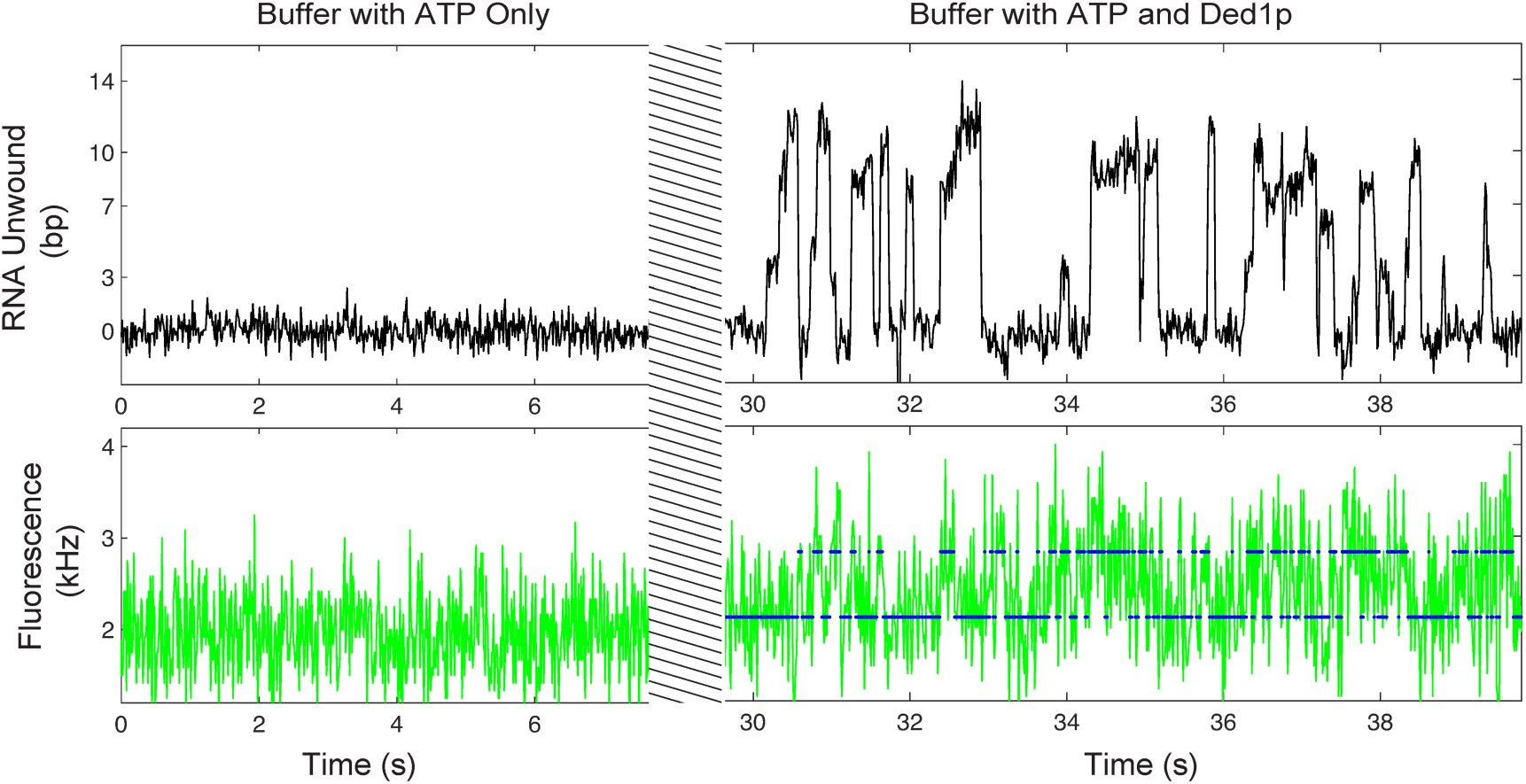
Example of simultaneous measurement by high-resolution tweezers (upper data) and fluorescence (lower data) showing RNA hairpin unwinding and rezipping when the hairpin tether is immersed in helicase (500 nM) and ATP (2.5 mM) containing solution (after t = 30 s) in comparison to stable, no-activity, control measurements in the ATP-only solution (before t = 30 s). The hatched bars cover the period when the tether is transferred between solutions (and data cannot be accurately acquired). The HMM fit (blue line) to the fluorescence data is shown.

**Supplementary Figure 5:**
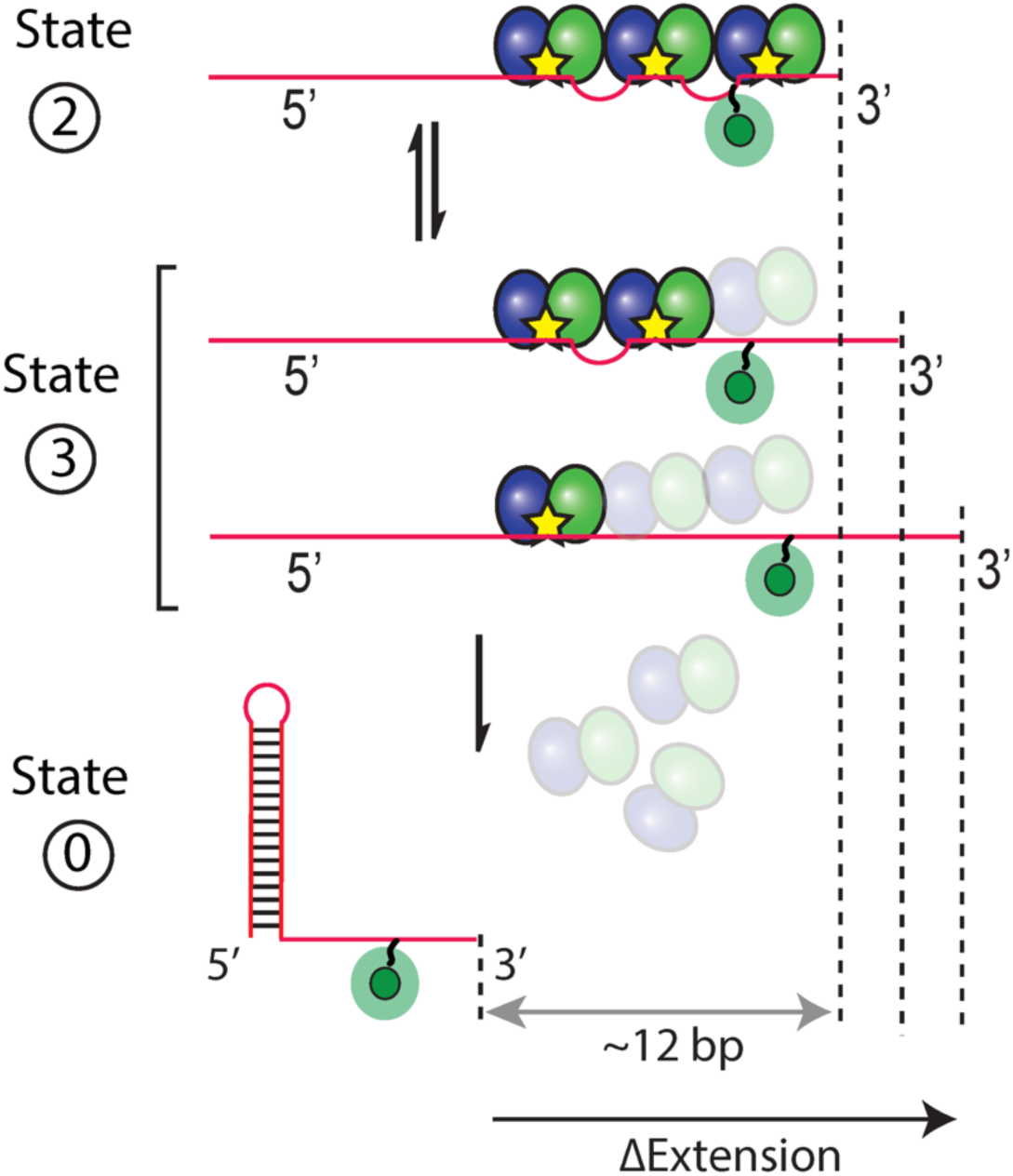
Cartoon of possible mechanisms of Ded1p dissociation from RNA backbone. In the unwound state 2, all three Ded1p promoters (blue and green oval balls representing Rec-A1 and A-2 domains) are bound to the RNA substrate (shown in red). The unwound states 3 and 3’ represent the partial dissociation of Ded1p promoters that could lead to complete dissociation or a situation where one or two proteins “hang on” while the last protein is still bound to the RNA, resulting in the relaxation of the RNA substrate. Complete dissociation of all three Ded1p promoters leads to the zipped state (0-state).

